# Linear-Nonlinear-Time-Warp-Poisson models of neural activity

**DOI:** 10.1101/194498

**Authors:** Patrick N. Lawlor, Matthew G. Perich, Lee E. Miller, Konrad P. Kording

## Abstract

Prominent models of spike trains assume only one source of variability – stochastic (Poisson) spiking – when stimuli and behavior are fixed. However, spike trains may also reflect variability due to internal processes such as planning. For example, we can plan a movement at one point in time and execute it at some arbitrary later time. Neurons involved in planning may thus share an underlying time-course that is not precisely locked to the actual movement. Here we combine the standard Linear-Nonlinear-Poisson (LNP) model with Dynamic Time Warping (DTW) to account for shared temporal variability. When applied to recordings from macaque premotor cortex, we find that time warping considerably improves predictions of neural activity. We suggest that such temporal variability is a widespread phenomenon in the brain which should be modeled.

## Introduction

A central goal in computational neuroscience is to build predictive models of neural activity and variability. Some variability comes from the stimuli and behaviors used in experiments (Hubel and Wiesel, 1962; Wurtz, 1969a,b). But even when experimental conditions are matched, neural activity can vary considerably (Reich et al, 1997; Steveninck et al, 1997; Kisley and Gerstein, 1999; Cai and Kass, 2005; Carandini, 2004; Lin et al, 2015). Some of this variability is due to the stochastic (Poisson) nature of spiking, but other potential sources exist. For example, neurons often share variability with their neighbors due to connectivity (Shenoy et al, 2013; Okun et al, 2015; Stevenson et al, 2012, 2008; Pillow et al, 2008) and common input (Vidne et al, 2012; Nordstrom et al, 1992). Crucially, internal brain states – such as attentional state – vary from trial to trial in spite of our best efforts to perform controlled experiments. At the neuronal level, this may mean that a population of neurons receives varying inputs on a trial-to-trial basis, leading to different activity patterns (Cohen and Maunsell, 2009; Mitchell et al, 2009; Rabinowitz et al, 2015). We are only beginning to appreciate these non-Poisson sources of variability.

One approach that has improved our understanding of trial-to-trial variability is the latent variable model (LVM) (Vidne et al, 2012; Yu et al, 2009; Goris et al, 2014; Chase et al, 2010; Lawhern et al, 2010; Pfau et al, 2013). LVMs attempt to model influences on neural activity that are difficult to measure or control directly. For example, one can model the effect of unobserved common input or gain modulation on a neural population (Vidne et al, 2012; Goris et al, 2014), leading to better model fits. Although typical LVMs can model considerable variability, it can be difficult to interpret the latent variables themselves since they don’t correspond to experimental variables.

Some types of neural activity can be dissociated from behavioral outputs. That is, there can be jitter between neural activity and events in the world such as movement or sensation (Kollmorgen and Hahnloser, 2014; Aldworth et al, 2011, 2005; Gollisch, 2006). For example, we can plan an arm movement at one point in time and subsequently choose the time of execution (Haith et al, 2016). This simple example reminds us that only primary input and output neurons need be closely coupled to the outside world. The retina most directly reflects the photons entering the eye, and spinal motoneurons reflect muscle activation. For brain areas in between, however, the temporal coupling to external factors could be much weaker; the time course of behavior-related neural activity in higher-level brain areas may instead vary from one trial to another. This temporal variability is likely to be shared across neurons that jointly solve a computational task such as generating behavior. Modeling such variability promises to lead to better models of neural activity.

Here we present a model for shared neural activity whose time course varies from trial to trial. We combine two well-known techniques: the Linear-Nonlinear-Poisson (LNP) model and Dynamic Time Warping (DTW). We label the corresponding variability “time-warp” variability. This model is appealing because it combines the interpretational simplicity of LNPs with population-level temporal dynamics. We first present a generative model for time-warp variability, which is an extension of the LNP. We develop a method for recovering the DTW and LNP parameters, and explore its behavior with simulations. We then apply our algorithm to simultaneously recorded neurons in macaque dorsal premotor cortex (PMd) during a reaching task. We find that accounting for time-warp variability improves model predictions, and explains an amount of variability comparable to spike history and reach-direction tuning.

## Results

Our research is partially motivated by anecdotal observations of shared temporal variability. Such variability can be seen in example data from a reaching task (Fig. 1). Four “stripes” of activity are visible, which correspond to four reaching movements. The time course of activity is quite similar across neurons, but varies from one reach to another reach. Both the temporal offset (relative to the onset of movement; Fig. 1, interval indicated by blue line) and the width of the stripe vary (Fig. 1, interval indicated by orange shading). It is this temporal variability that we seek to model in this paper.

**Figure 1:**
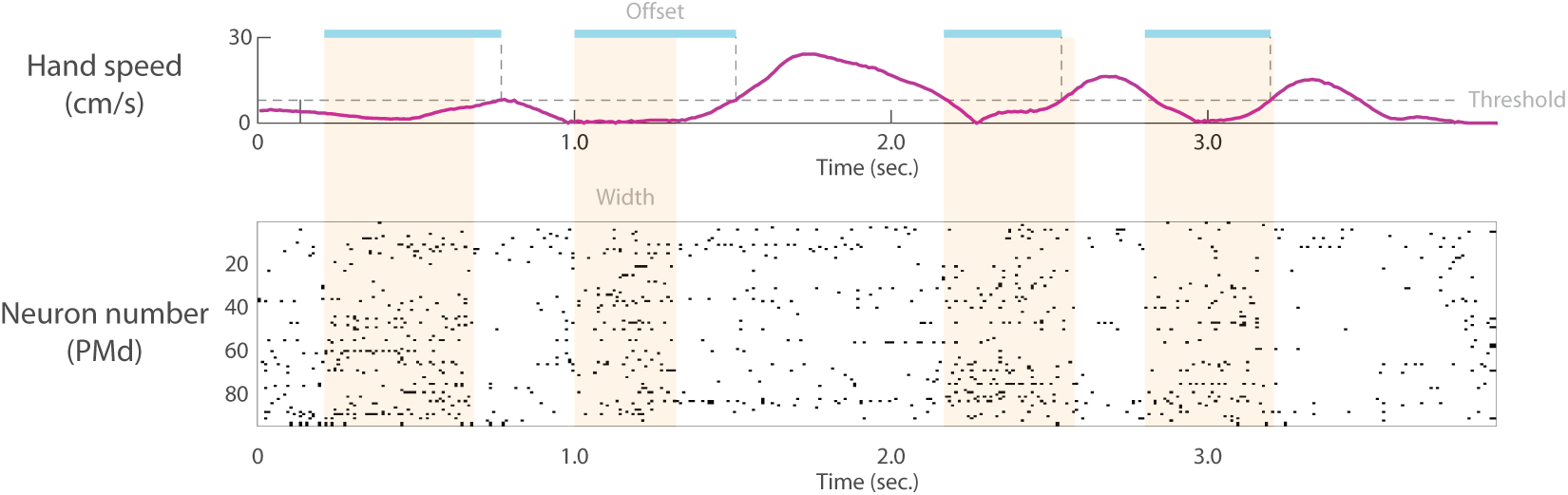
Single trial time-warp variability in macaque PMd. The monkey makes a sequence of four reaches to targets. The hand speed is shown in the top panel. The bottom panel shows a spike raster of 95 simultaneously recorded PMd neurons. Four stripes of activity corresponding to four reaches are visible by eye (orange shading, drawn by hand to approximately enclose stripe of activity). Both the temporal offset of these stripes with respect to movement onset (blue shading, drawn by hand from onset of activity to time of movement onset defined by speed threshold) and width of the stripes (orange shading) vary from reach to reach. Reaches are not matched for direction and other kinematic parameters (kinematics is considered in more depth in the Results section). The neural data here simply serves to provide intuition for time-warp variability.

### Generative Model

#### The standard Linear-Nonlinear-Poisson Model (Poisson Generalized Linear Model)

Here we extend the Linear-Nonlinear-Poisson (LNP) framework, a prominent model of neural activity, to accommodate time-warp variability. The LNP model essentially generalizes multiple linear regression to spike (Poisson) data, making it possible to ask what factors predict spiking. We begin with the generative process of the LNP (Fig. 2). First, explanatory covariates such as reach direction, 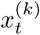 are linearly combined with scalar weights, *θ*_*k*_. Here, 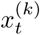 is the value of covariate *k* at time *t*, in a time bin of size *dt*; *x*^(*k*)^ is a *T* × 1 vector representing the time series of covariate *k*; *X* is a *T* × *K* matrix representing the collection of all *K* covariate time series; *θ*_*k*_ is a scalar weight corresponding to covariate *k*; Θ is a *K* × 1 vector of weights.

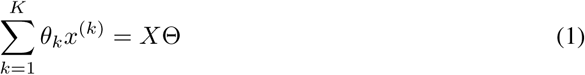

**Figure 2:**
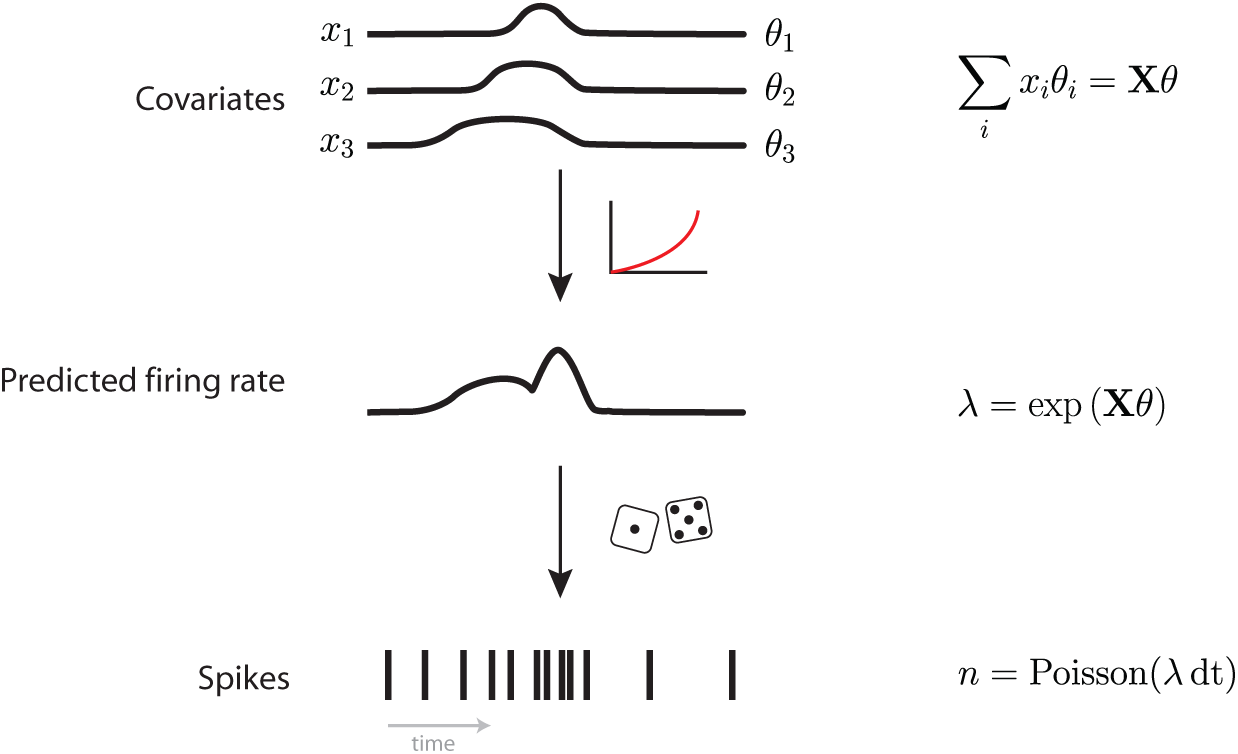
Linear-Nonlinear-Poisson model of neural activity. The LNP model. Explanatory covariates are linearly combined and passed through a nonlinear function to yield a non-negative firing rate. Spike counts are then viewed as Poisson noise around this underlying firing rate.

Next, these weighted covariates are passed through a nonlinear function, typically exp(). The exponential function is the canonical inverse link function of the Poisson Generalized Linear Model (GLM) (Nelder and Baker, 1972), so called because it links the covariates (positive-or negative-valued) with the domain of the Poisson distribution (non-negative-valued). This step yields a time series of the mean firing rate, λ which is a *T* × 1 array.

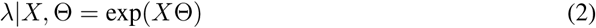

Spike count *n*_*spikes*_ in each time bin of duration *dt* is then modeled as Poisson given this firing rate.

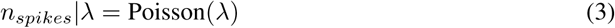

This leads to the following likelihood function for a single spike train:

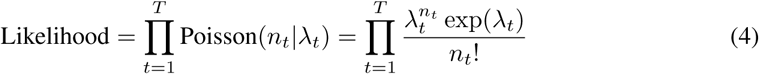

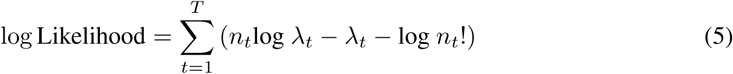

Data points are written as conditionally independent given the covariates, *X*. Spiking history, for example, can be accounted for as a covariate in *X*.

It is important to note that this type of model is typically fit *across trials* (or across reaches in Fig. 1). Using only regressors that relate to the nature and timing of external variables, e.g. sensory stimuli or movement kinematics, this model cannot account for internal trial-by-trial variability. In other words, the model will approximate the average response of the neuron to features in the outside world. This is the property of the LNP which we aim to improve.

But how can we model activity that differs in time course from trial to trial but that doesn’t depend on stimuli or behavior? Our intuition is that neural activities during one trial may start earlier or later than during another trial, or last longer or shorter. More generally, time may be nonlinearly warped between population neural activity and variables of the outside world. Although it may be different on each trial, we hypothesize that it is shared across neurons and can be modeled.

### Dynamic Time Warping

There exists an algorithm that addresses such timing problems: Dynamic Time Warping (DTW) (Sakoe and Chiba, 1978). DTW has been used in a variety of applications (Berndt and Clifford, 1994), including neuroscience (Reich et al, 1997; Perez et al, 2013; Victor, 2005; Victor and Purpura, 1996), to understand how two time-varying signals relate to one another. More specifically, DTW finds the optimal transformation to align one signal and its counterpart when the signals differ by a time warp. Intuitively, for every time bin in signal 1, the algorithm will find the time bin in signal 2 with the best correspondence. If both signals are of length *T*, this requires *T*^2^ comparisons. It is convenient to view these *T*^2^ comparisons as a matrix (Fig. 3) which measures the compatibility between the two signals for each possible alignment.

**Figure 3:**
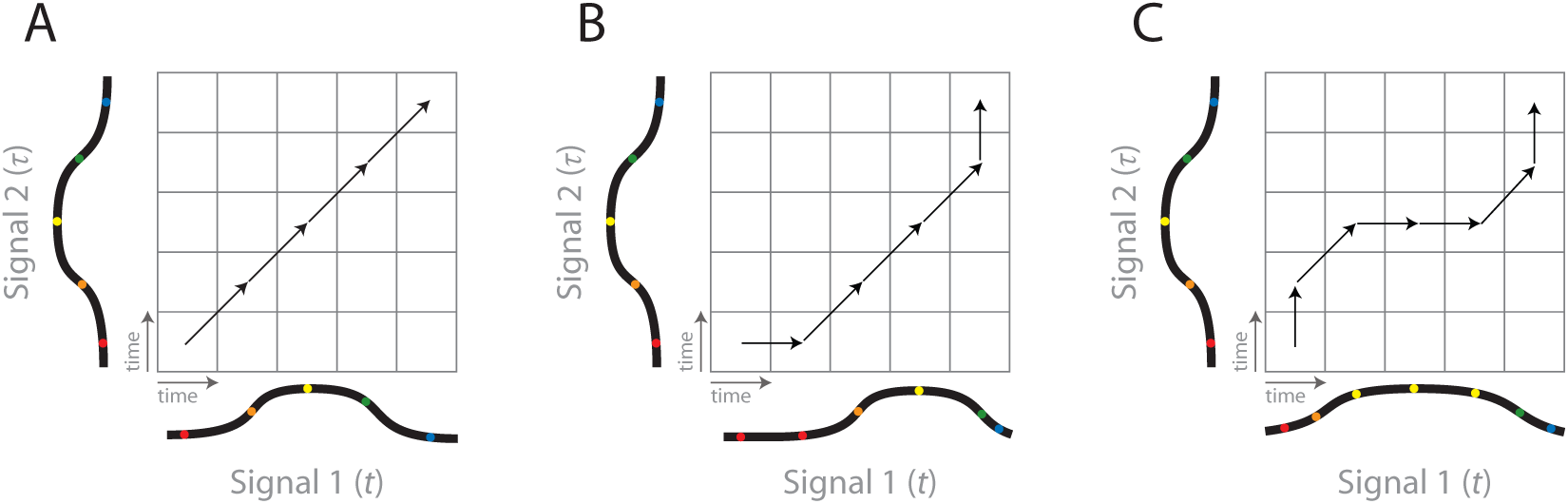
Time warping schematic. **A**) A simple application of the DTW algorithm to find the best alignment between two signals. In this case, the signals are well aligned and do not require shifting or scaling. **B**) In this case, however, one signal lags the other, and a transformation is required to align them. These transformations are expressed as “steps” taken through the matrix (because the signals are discrete). Diagonal steps correspond to no shifting or scaling. Any other type of step will lead to a shift or scale operation. **C**) In this case, signal 1 is stretched relative to signal 2, requiring a scaling operation to align them.

The goal of the DTW algorithm then becomes to find the best path through this *T* × *T* matrix that begins with both signals at *t* = 1 and ends with both signals at *t* = *T*. Because the two signals are discrete, this path will take the form of steps (between time bins). In the simplest case, individual steps can be horizontal, vertical, or diagonal (Fig. 3B). Diagonal steps (up 1 and over 1) do not produce a warp because time moves forward at the same rate in both signals. A horizontal step introduces a delay in Signal 2 relative to Signal 1 (Fig. 3B). Similarly, a vertical step introduces a delay in Signal 1 relative to Signal 2. By delaying time in one signal relative to the other, non-diagonal steps can effectively produce either an offset or scaling operation in time – a time warp (offset and scaling operations are conceptually useful for understanding the algorithm, but are not explicitly modeled beyond allowing non-diagonal steps). The full path is a series of these steps, and can therefore produce a complex combination of multiple offset and scaling operations.

Further, DTW constrains that time must not move backwards in either signal. The path through this matrix must also be continuous: any step of the path must be possible given the previous step. Although allowed steps most often include horizontal (over 1), vertical (up 1), and diagonal (up 1 and over 1), other types may be desirable for a given application (e.g., over 1, up 2; Fig. 4 and Methods section). In addition, it is possible to specify a prior distribution over these steps. This may be desirable if some types of steps are a priori more likely, or to avoid overfitting noisy signals. Other than a uniform prior over types of steps, the most commonly-used prior expresses a preference for diagonal steps.

**Figure 4:**
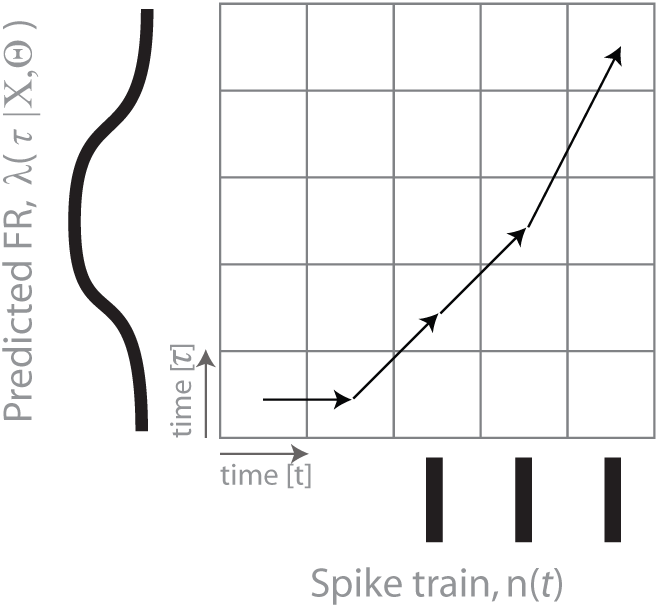
LNP+DTW Schematic. In the combined model, we use DTW to align the trial-averaged LNP model prediction with the spikes measured in a particular single trial.

More formally, the DTW algorithm optimizes the following cost function (Berndt and Clifford, 1994; Sakoe and Chiba, 1978):

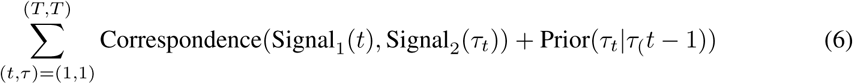

The sum is taken over the entire DTW path, for each time bin *t* of *T* in signal 1, and each time bin *τ* of *T* in signal 2. We will summarize the DTW path with the parameter **t** = {*t*_1_,*t*_2_,…,*t*_*T*_} and *τ* = {*τ*_1_, *τ*_2_,…, *τ*_*T*_}. Together they specify the coordinates (*t*, *τ*) of the path in the alignment matrix. ‘Correspondence()’ is some metric of agreement (or distance) between signal 1 and signal 2. The global optimum can be found using dynamic programming (see Supplemental Methods for intuition) (Berndt and Clifford, 1994; Sakoe and Chiba, 1978). Here, we extend DTW to accommodate vector-valued signals by choosing the Correspondence() function appropriately (see Methods). We will make use of this in the next section in order to use DTW with simultaneously recorded spike trains.

#### Combining Linear-Nonlinear-Poisson and Dynamic Time Warping approaches

Now we seek a way to combine the LNP model with DTW into a statistical model. The goal is to use the LNP model to capture the trial-averaged responses of neurons, and then to use DTW to correct the LNP model on a trial-by-trial basis. Intuitively, we accomplish this by choosing the DTW input signals appropriately (Fig. 4): signal 1 is the measured spike train for a single trial, and signal 2 is the LNP model prediction for that trial. This allows us to find the best way to transform the LNP model prediction —- which is effectively trial averaged —- to best agree with the spikes measured in a particular trial.

More formally, we construct a statistical model that specifies the relevant variables, how they relate to one another, and their distributions (Fig. 5). We are ultimately interested in modeling the number of spikes 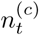 for cell *c* that occur at time *t* in a time bin of duration *dt*. We assume that 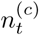 is Poisson-distributed with a mean given by 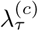, which varies in time (a binned inhomogeneous Poisson process). 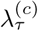, in turn, depends on the parameters of the standard LNP process, Θ^(*c*)^, as well as a time-warp path, *τ*. The LNP parameters, Θ^(*c*)^, describe a neuron’s dependence on variables (e.g., reach direction), *X*. Each neuron *c* is described by its own set of parameters Θ^(*c*)^, which are assumed to be fixed across trials and independent of other cells. The time-warp parameter, *τ* = {*τ*_1_, *τ*_2_,…,*τ*_*T*_}, is modeled as a nonlinear mapping between the LNP model time, *τ*, and spike time, *t*. Together, *τ* and *t* specify the location of the path in the alignment matrix that relates the value of the LNP model at time *τ* and the spike train at time *t* (we will use *τ*_*t*_ as shorthand for the (*t*, *τ*) ordered pair). We assume that *τ* is shared across neurons, but varies from trial to trial.

**Figure 5:**
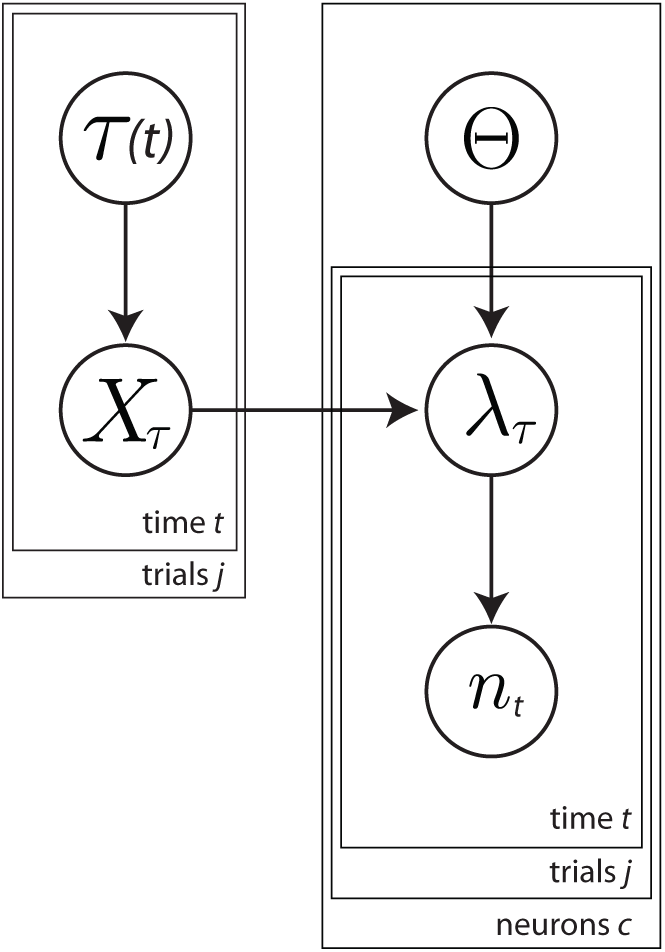
Generative model for time-warp variability. A given neuron’s spiking is a function of the mean firing rate, λ, as in the standard Linear Nonlinear Poisson model. The LNP parameters of each neuron, Θ, are fixed across all trials. Spiking is also influenced by a trial-specific time-warp, *τ* which is shared across neurons. For simplicity, we have omitted the superscripts identifying the parameters for each neuron and trial.

To fit this model to data, we seek the values of Θ and *τ* that are maximally likely given the data; i.e., we seek to maximize P(Θ, *τ*|*n*). This expression is difficult to optimize directly, so we fit these parameters using an alternating approach similar to Expectation-Maximization (Dempster et al, 1977). We fix one set of parameters while fitting the other, and then switch, alternating until convergence (see Methods and Supplemental Methods for full description). For the LNP portion of the model (for a single neuron and a single trial), the parameters are given by

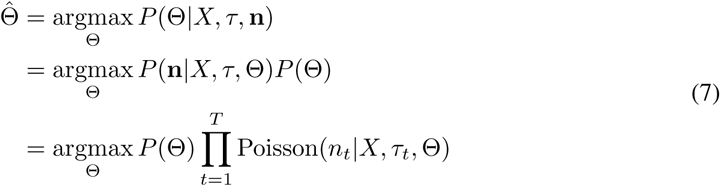

This is a Poisson Generalized Linear Model which can be optimized using standard methods (the cost function is convex, so the global optimum can be found with gradient ascent). For the DTW portion of the model, the parameters are given by

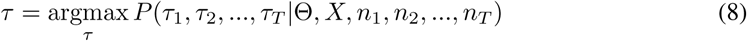

Which can be written in the same form as the cost function of DTW (Eqn. 6, see Methods).

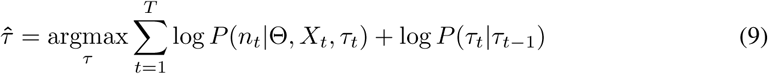

Like the DTW cost function, the first term represents the compatibility between the two signals (the measured spikes and the time-warped LNP model predictions). The natural choice for this term is the Poisson likelihood function

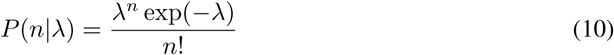

Where λ = exp(*X*Θ). The second term represents a prior distribution over what types of steps are possible and how likely they are. This ultimately leads to the following cost function (see Supplemental Methods).

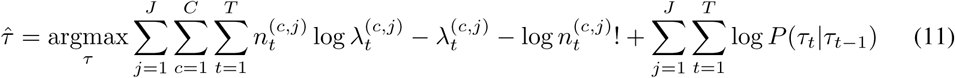

The sums are taken over *C* cells, *J* trials, and *T* time bins per trial. Spike time is indexed by *t*, and LNP model time is indexed by *τ*. This model can be fit to data and tested using cross-validation (see Methods).

### Simulations

In order to test whether our model could correctly capture time-warp variability, we simulated data with known parameters and attempted to recover those parameters. We first simulated a population of PMd neurons using an LNP model of movement-related activity. We then added shared time-warp variability on a trial-by-trial basis (see Supplementary Methods for procedure). We were able to accurately recover 1) the receptive field parameters for individual neurons (preferred direction, temporal response properties; Fig. S2), and 2) the time-warp parameters which characterize the shared temporal variability (see Supplementary Results, Fig S1). These simulations demonstrated that our model behaved as expected, and would not give spurious results when applied to real data.

#### PMd recordings from monkeys during a reaching task

We next applied our model to real neural data in order to quantify the magnitude of time-warp variability. We analyzed extracellular recordings from PMd during a random-target reaching task (Fig. 6). In this task, monkeys controlled an on-screen cursor by making reaching movements using a planar manipulandum and were rewarded for successfully moving to displayed targets. In contrast to center-out reaching tasks with a single target per trial, this task used a sequence of four targets per trial. These targets appeared sequentially in any location within the workspace and the monkey was rewarded after four successful target acquisitions. There were minimal time restrictions (see Methods), meaning that the monkeys made series of smooth, freely-initiated movements. We used this task because it is somewhat more naturalistic than a center-out reaching task and spanned a broader range of positions and velocities. As a result, we expected less-stereotyped movement planning and execution.

**Figure 6:**
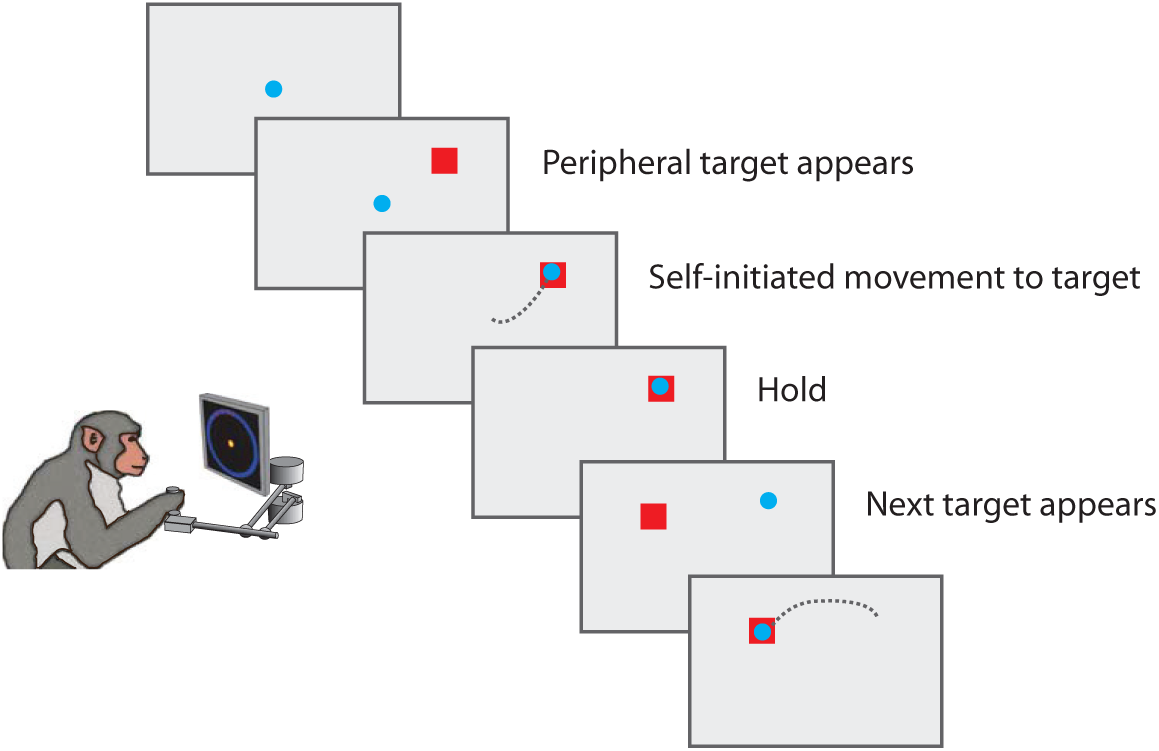
Reaching experiment schematic. In this task, the monkey controlled a cursor using a manipulandum. A visual target indicated the reach target, and the monkey was free to initiate movement. After reaching the target and pausing for a brief hold period, another target would appear in a pseudo-random location. Four targets were shown per trial.

The task was performed by two macaque monkeys with electrode arrays chronically implanted in PMd. Using standard spike sorting techniques, we identified 95 neurons in Monkey M and 49 neurons in Monkey T.

To assess model performance, we constructed two LNP+DTW models: 1) a basic model to gain intuition about time-warp variability, and 2) a comprehensive model to compare time-warp variability with other known sources of variability.

For the basic LNP model, we analyzed neural activity relative to movement onset (as measured by a speed threshold crossing). This model allowed for reach-direction tuning, variable temporal responses, as well as a spatially untuned response (i.e., activity locked to movement onset but independent of reach direction (Fernandes et al, 2013); see Methods for details and Fig. 11 for graphical schematic of model). Note that we did not include model components related to target presentation time as this was not precisely recorded during the experiments (see Methods and Discussion). We then fit the LNP+DTW model (i.e., including time warping) as described in the Methods section.

We first asked whether accounting for time-warp variability improved model predictions at the single-trial level. We therefore visually examined model predictions before and after the time-warp correction, and compared them with the measured spiking activity. For many reaches, the uncorrected model predictions did not correspond well to the measured spiking activity (Fig. 7). The actual spiking activity sometimes preceded or lagged the uncorrected model predictions (Fig. 7A). In other cases, bouts of spiking activity were longer lasting (Fig. 7B) or briefer (Fig. 7C) than predicted by the uncorrected model. After correcting for time-warp variability, however, model predictions better matched the measured activity. By eye, a model that accounts for time warp makes better predictions of spiking activity than one that does not.

**Figure 7:**
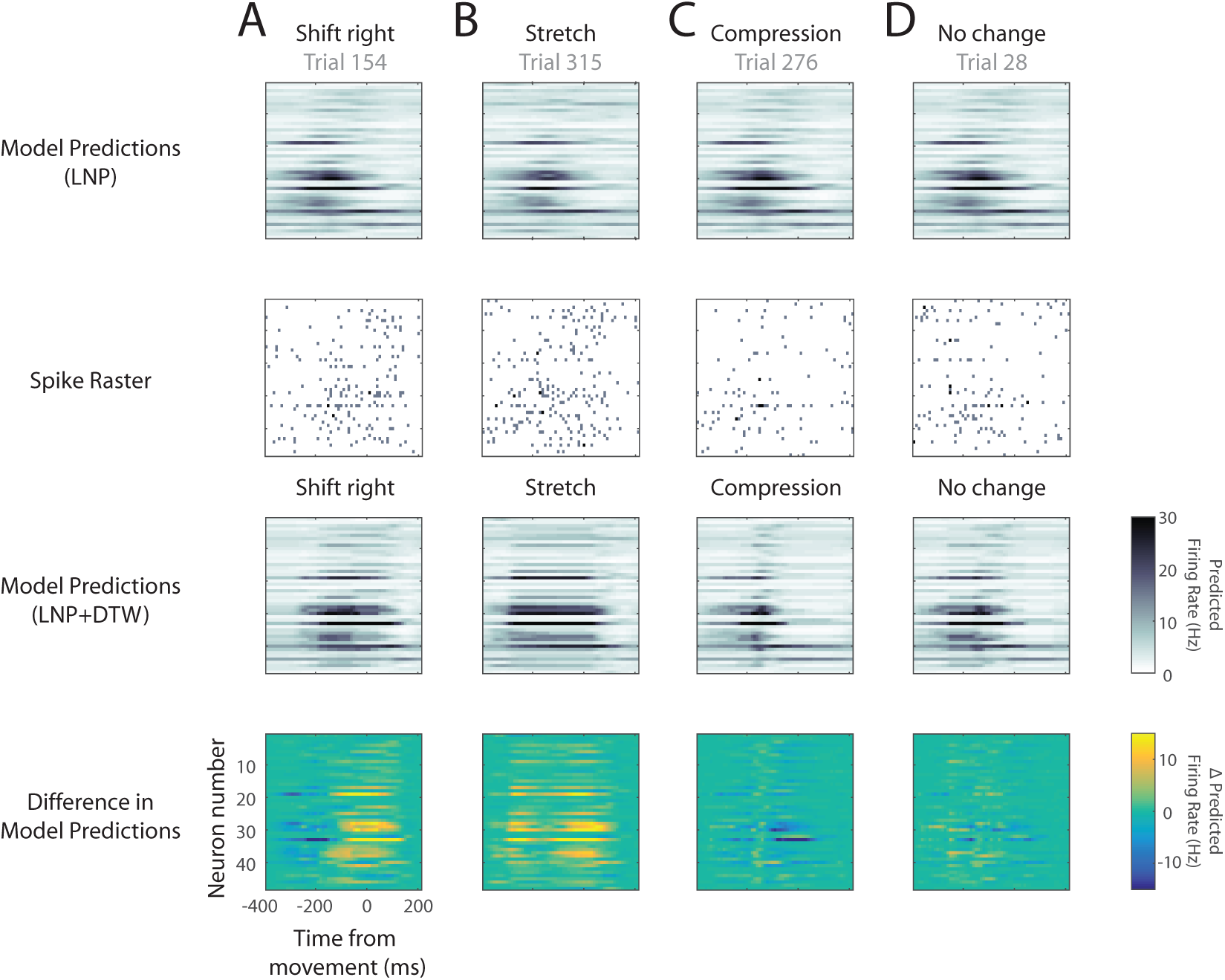
Single-trial corrections for time-warp variability. Spike rasters (second row) and predicted firing rates (first and third rows) for 48 simultaneously recorded PMd neurons from Monkey M. Within a raster, each row corresponds to a different neuron. Each large column (A-D) corresponds to a different example reach. To simplify comparisons across the displayed reaches, all four were chosen from the same 45 degree reach direction. As a result, the LNP predictions look similar for each of the four (first row). Note that the LNP predictions appear to be smooth because they reflect the “underlying firing rate”. Spikes are viewed as Poisson noise around these smoothly varying functions. **A**) In this trial, the spiking activity (second row) lagged the LNP model predictions (first row). The LNP+DTW model thus shifted the predictions to be later in time (third row). The fourth row shows the differences in predictions due to time-warp correction. **B**) In this trial, the bout of spiking activity (second row) lasted longer than predicted by the LNP model (first row). The LNP+DTW model thus stretched the predictions (third row). **C**) In this trial the bout of spiking activity was briefer than predicted by the LNP model (first row). The LNP+DTW model compressed the predictions (third row). **D**) In this trial, the LNP prediction (first row) was similar to the spiking activity (second row), and little correction was needed.

We also examined DTW alignment paths at the single-trial level to understand how the algorithm accomplished the transformations we observed. To do this, we plotted the inferred DTW path alongside the LNP model predictions and spike trains (Fig. 8). This allowed us to visualize the correspondence between the spike trains at time *t* and model predictions at time *τ* for each point (*t*, *τ*) along with DTW path. We also colored the alignment matrix at every point according to this correspondence (i.e., according to the Poisson log likelihood evaluated at (*t*, *τ*)). As expected, the DTW path in each trial traverses the high-correspondence regions in the alignment matrix, thereby maximizing the path’s log likelihood. The shape of the these DTW paths can be understood intuitively by comparing them with the simpler examples in Fig. 3. In trial 154, the spikes exhibit a “shift” to the right, which the DTW path accomplishes by “skipping” time (many horizontal steps) at the beginning of the trial (Fig. 8A, Fig. 3B). In trial 315, the spikes exhibit a “stretch” (Fig. 8B, Fig. 3C). And in trial 28, in which little transformation is needed, the DTW path is relatively diagonal (Fig. 8D, Fig. 3A). Thus, the we found that the inferred DTW paths made sense conceptually given the transformations they produced.

**Figure 8:**
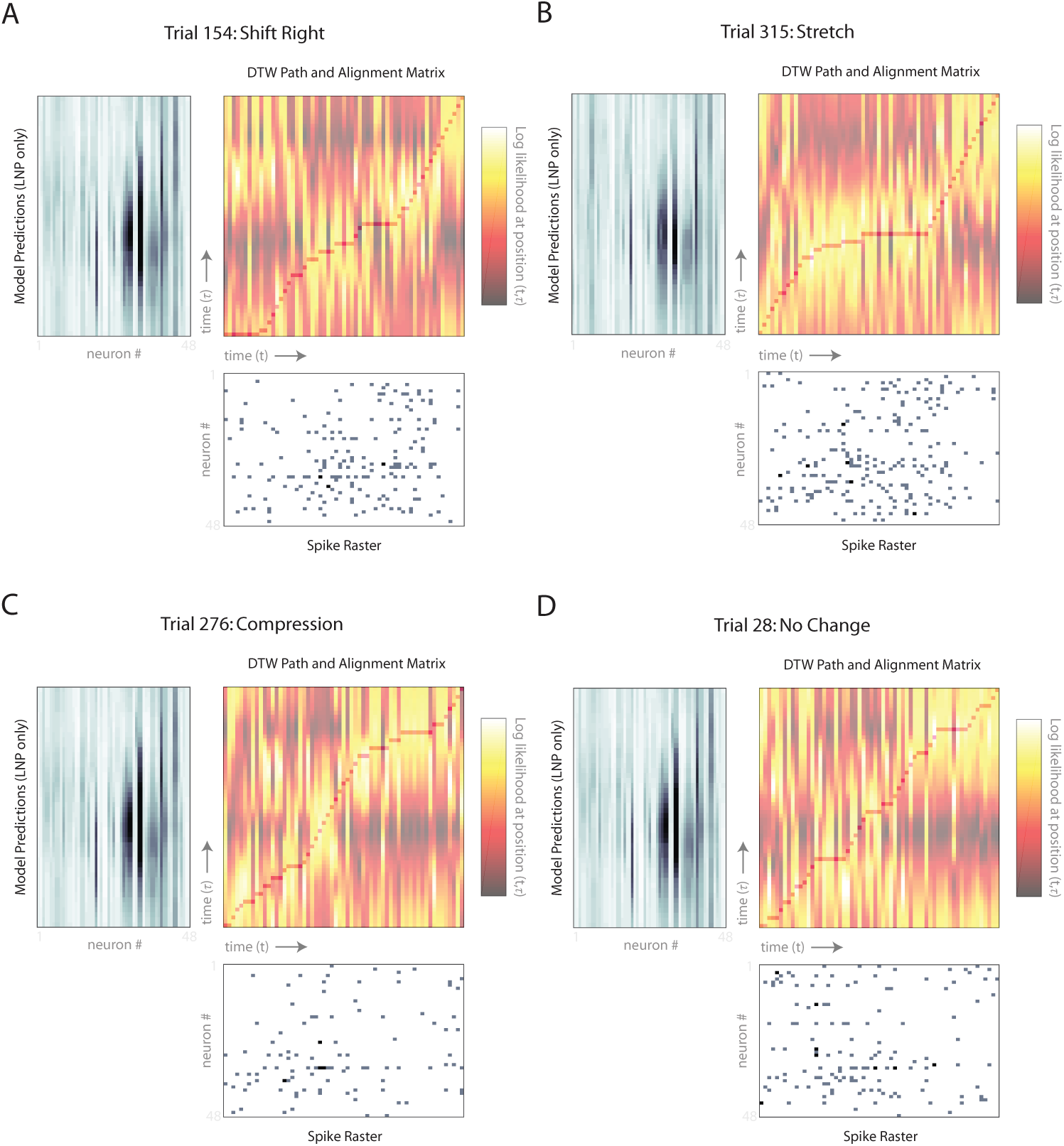
Single-trial warp paths. Warp paths for the four trials shown in Fig. 7 A-D which correspond to panels A-D here. Each panel contains three subpanels: LNP model predictions for all 48 neurons (top left subpanel, rotated 90 degrees), spike raster for all 48 neurons (bottom right subpanel), and the alignment matrix with warp path (top right subpanel). This plot can be read like Figs. 3 and 4 with the exception that there are multiple neurons on each time axis rather than a single neuron. The alignment matrix is colored by how well position *t* in the spike raster corresponds to position *τ* in the LNP model predictions in terms of Poisson log likelihood (NB: we z-scored the alignment matrix columns for visualization purposes). Superimposed on the alignment matrix is the inferred warp/alignment path that best matches the LNP model predictions with the spike raster.

Next, we asked whether correcting for time warp led to improved predictive power. We found in Monkey M that for approximately half of the neurons modeled (28/48), time warping significantly improved predictive power (at *p* < .05, bootstrap test) (Fig. 9, 10 left panel). Further, we found that the median effect size of time-warp variability across the population was considerably greater than zero (Fig. 10 right panel, basic model, red markers). Accounting for time-warp variability thus improves predictive power beyond that of a basic LNP model.

**Figure 9:**
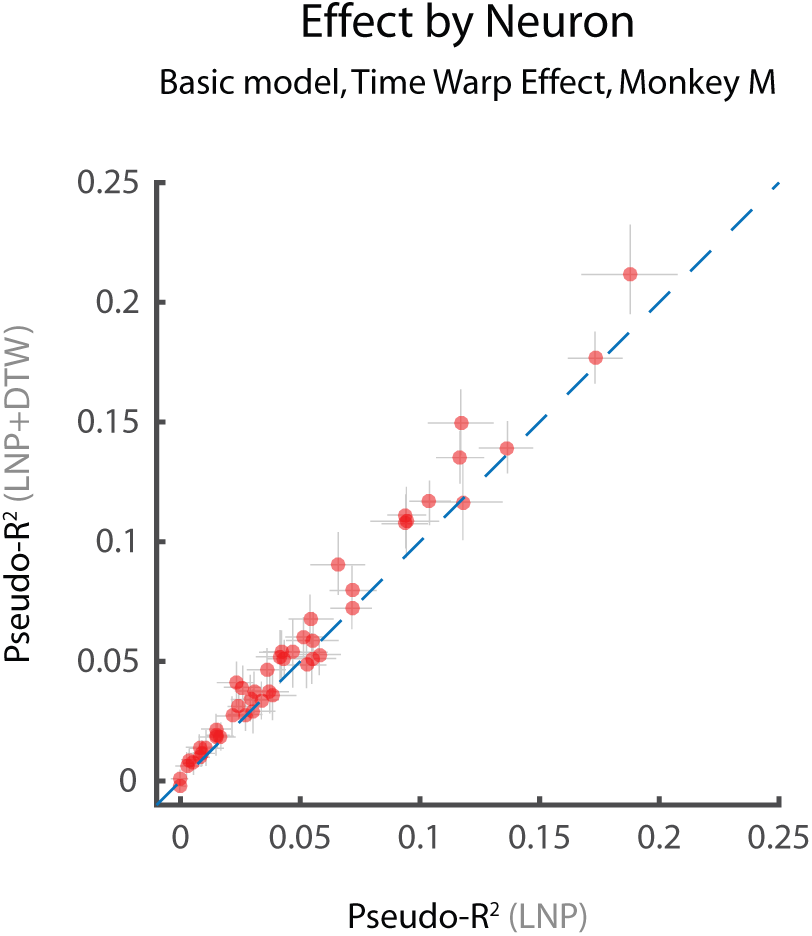
Summary statistics for time-warp variability: Pseudo-*R*^2^. Predictive performance of LNP+DTW models of neural activity in terms of Pseudo-*R*^2^. All performance metrics were computed using cross validation (see Methods). **A**) Scatterplot of Pseudo-*R*^2^ values before (LNP only) and after including time-warp variability (LNP+DTW). Each data point represents a single neuron. Points above the unity line indicate that a model including time-warp variability is better than one without it. Error bars represent bootstrapped 95% confidence intervals. Data shown for Monkey M (MM) only.

**Figure 10:**
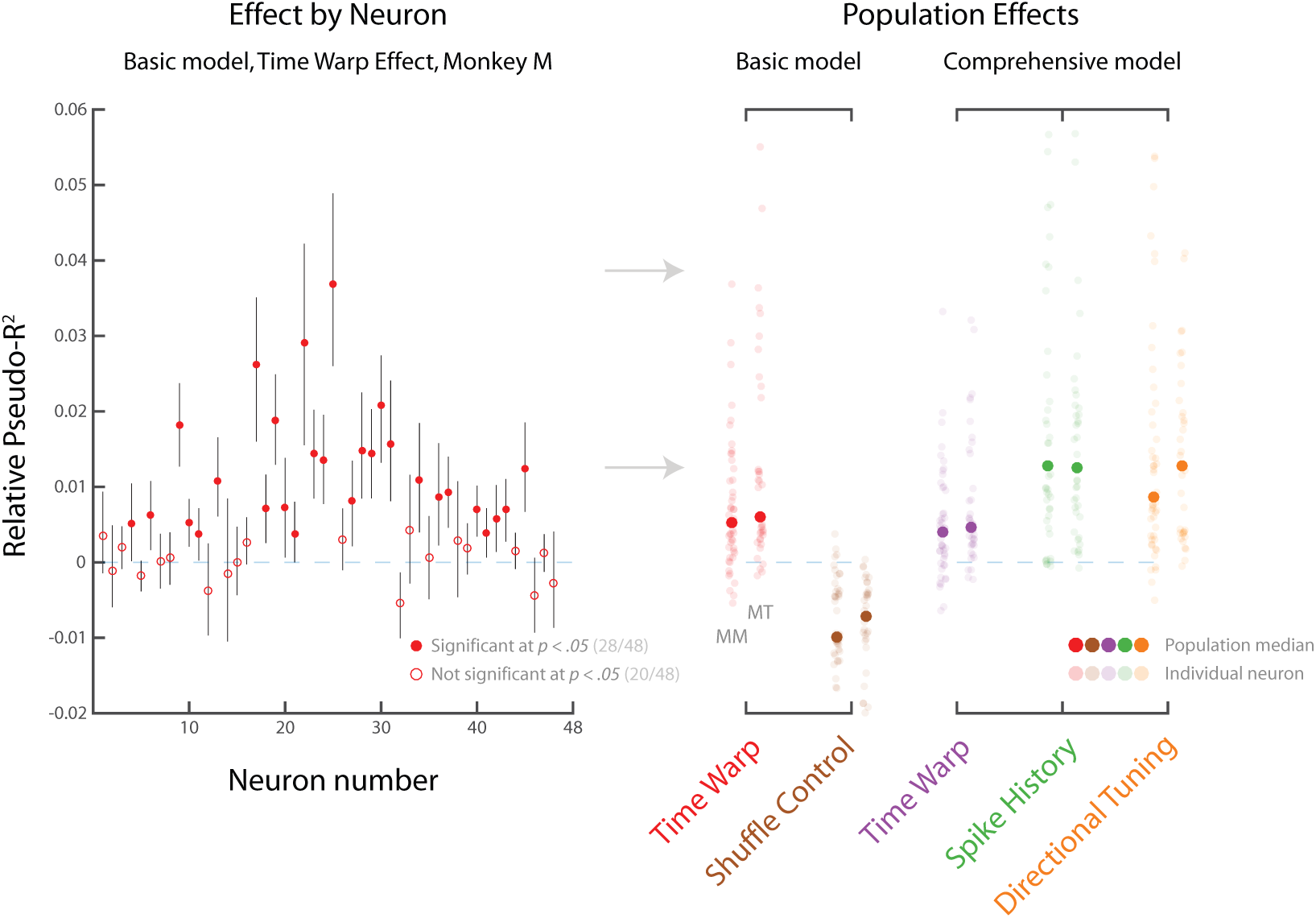
Summary statistics for time-warp variability: Relative Pseudo-*R*^2^. Effect sizes measured by Relative Pseudo-*R*^2^, a related measure of predictive accuracy that compares two models. Positive values indicate improvement, a value of zero indicates no change, and negative values indicate worsening. Error bars represent bootstrapped 95% confidence intervals. Left panel: In the basic LNP+DTW model, 28/48 individual neurons were predicted better by the LNP+DTW model compared to the LNP alone for MM (left panel). Right panel: Median effect sizes (across neurons) for two different models: the basic LNP+DTW model, and the comprehensive LNP+DTW model. Effect sizes represent marginal change in predictive power between the model with all components, and a model leaving out one component. In the basic LNP+DTW model, the median effect size of time-warp variability is considerably greater than zero (red markers). A shuffle control leads to worsened predictions (brown markers, see text). In the comprehensive LNP+DTW model, we additionally accounted for reach-specific kinematics and spike history. The median effect size of time-warp variability remains greater than zero (purple markers), and is comparable in magnitude to spike history (green markers) and directional tuning (orange markers). Data shown for both Monkey M (MM) and Monkey T (MT).

**Figure 11:**
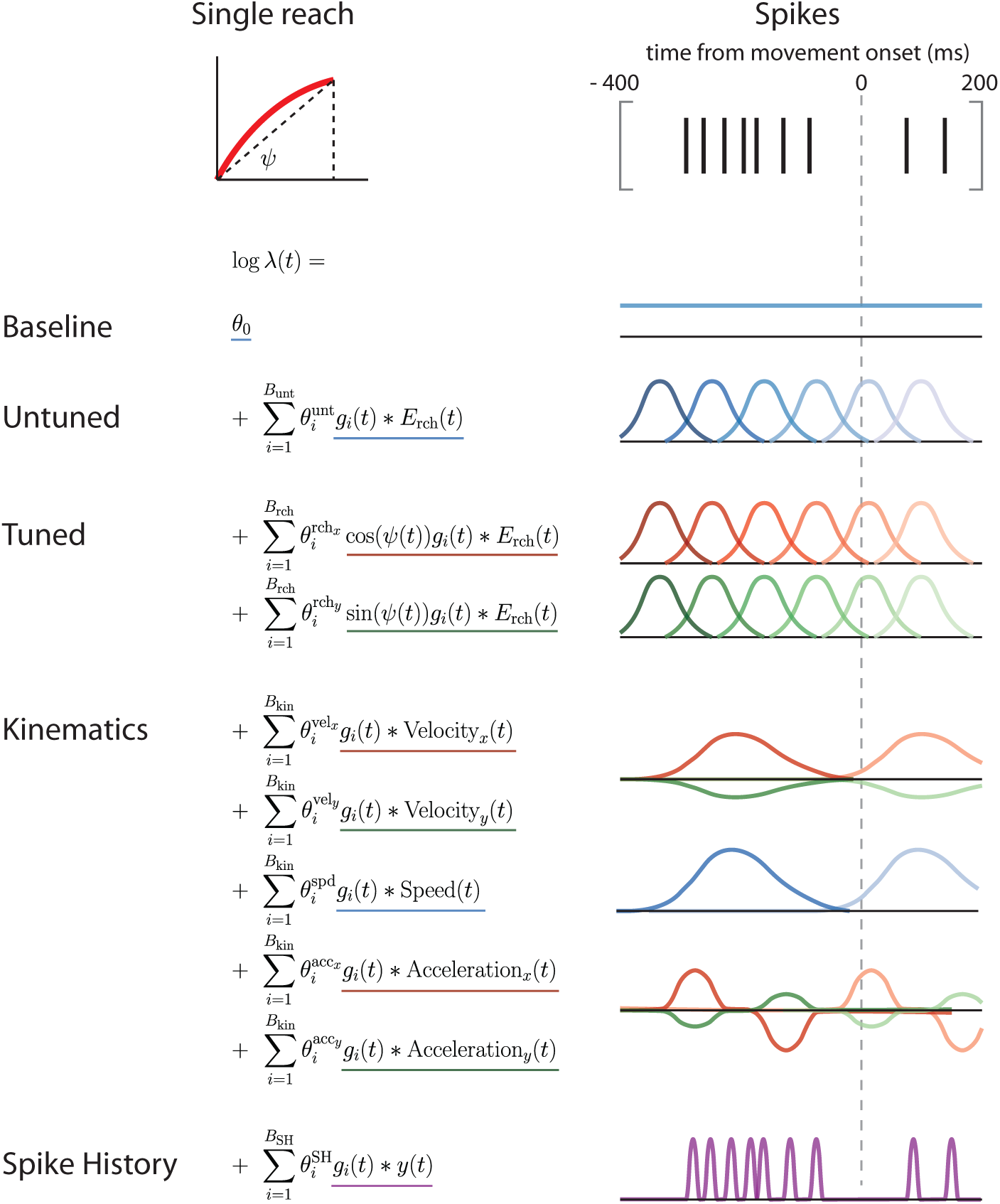
Comprehensive LNP model of neural activity. We model the spiking activity in a fixed window surrounding each reach (400 ms before, 200 ms after; 10 ms bins). The comprehensive model includes the following components: baseline firing rate; an untuned temporal response to each reach; a spatially tuned temporal response to each reach which depends on the direction, but not on the kinematics; components that depend directly on reach velocity and acceleration; and spike history terms.

It is possible, however, that accounting for time-warp variability simply improved the estimates of neurons’ average temporal responses, rather than improving estimates of trial-to-trial variability. Although the LNP model allowed for considerable temporal variability, it might still not describe all possible temporal responses adequately. For example, DTW could be performing some kind of generic operation that better shapes the temporal aspects of the LNP model predictions (e.g., by smoothing). If this were true, the calculated time-warp path should be similar across trials, and thus interchangeable across trials. To test this, we performed a shuffle analysis in which time-warp paths were calculated on the correct trials, but then applied to other trials at random (see Methods). We found that shuffling trials in this way led to predictions that were worse than those without time warping (Fig. 10, right panel, basic model, brown markers). This suggests that time-warp variability is unique to each trial, and that improvements in model predictions do not result from better modeling of the across-trial average neural responses.

Next, after having explored the behavior of the basic LNP+DTW model, we built a more comprehensive model. This model was designed to compare the importance of time-warp variability with other known sources of trial-to-trial variability. The comprehensive model’s LNP portion included the same covariates as the basic model (reach-direction tuning, variable temporal responses, spatially untuned response), as well as: instantaneous velocity, instantaneous speed, instantaneous acceleration, and spike history (see Methods for full model description). To compare the importance of each model component, we computed the predictive power lost by leaving it out. We found that the effect size of time-warp variability was smaller than, but of approximately the same order as, both spike history and directional tuning (Table 1; Fig. 10, right panel, comprehensive model). It is unsurprising that the effect is smaller than directional tuning, as movement planning is a primary function of PMd (Cisek and Kalaska, 2004). Importantly, the median effect size of time-warp variability was only modestly lower in the comprehensive model compared with the basic model (Fig. 10; Table 1, see Methods), suggesting that time-warp variability is largely a separate source of variability. Time-warp variability is therefore a relatively strong effect compared to other sources of variability in spike trains.

**Table 1:**
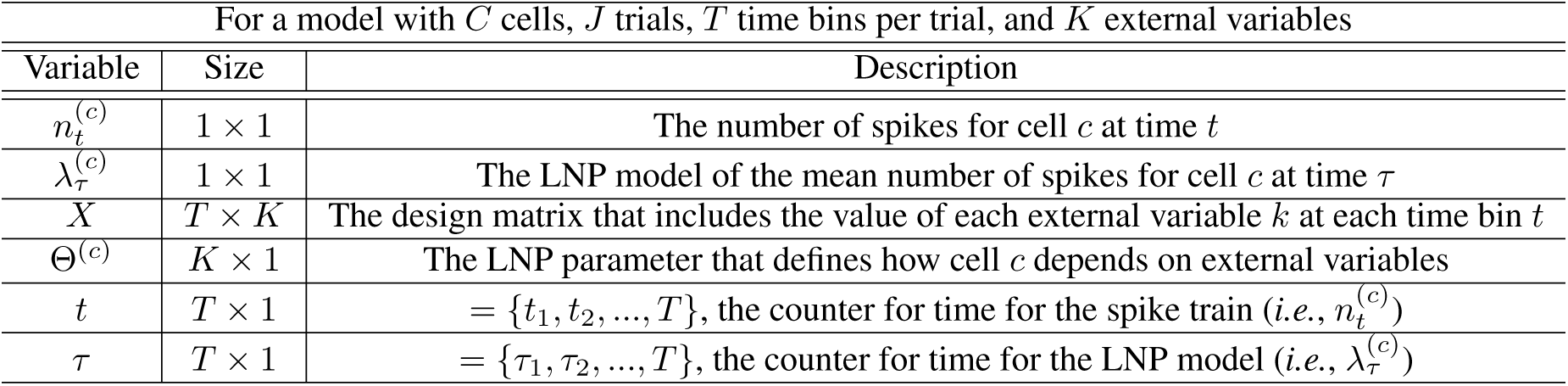
Definitions of model variables

| For a model with $C$ cells, $J$ trials, $T$ time bins per trial, and $K$ external variables | | |
| --- | --- | --- |
| Variable | Size | Description |
| $n_t^{(c)}$ | $1 \times 1$ | The number of spikes for cell $c$ at time $t$ |
| $\lambda_\tau^{(c)}$ | $1 \times 1$ | The LNP model of the mean number of spikes for cell $c$ at time $\tau$ |
| $X$ | $T \times K$ | The design matrix that includes the value of each external variable $k$ at each time bin $t$ |
| $\Theta^{(c)}$ | $K \times 1$ | The LNP parameter that defines how cell $c$ depends on external variables |
| $t$ | $T \times 1$ | $= \{t_1, t_2, \dots, T\}$, the counter for time for the spike train (*i.e.*, $n_t^{(c)}$) |
| $\tau$ | $T \times 1$ | $= \{\tau_1, \tau_2, \dots, T\}$, the counter for time for the LNP model (*i.e.*, $\lambda_\tau^{(c)}$) |

## Discussion

In this paper, we have developed a model for time-course variability that differs from trial to trial and is shared across neurons: time-warp variability. This model is notable for combining the simplicity of the single-neuron LNP with population-level temporal coupling. We have presented a generative model for this process, tested the inference algorithm with simulations, and applied it to real data from the macaque premotor cortex. We find that the effect size of time warping is roughly comparable to other well-appreciated sources of variability, including directional tuning and spike history.

Previous work in neuroscience has used ideas related to time warping to explore different aspects of temporal variability (Aldworth et al, 2011, 2005; Gollisch, 2006). Perez et al (2013) asked whether individual neurons better align to stimulus timing or motor timing in premotor areas. To do this, they used a model in which neural activity may either be a function of time relative to the stimulus, or of rescaled time relative to the movement. Victor et al (Victor, 2005; Victor and Purpura, 1996) have used the DTW algorithm to assess spike train similarity. Lakshmanan et al examines neuron-specific delays in the nervous system, and proposes a LVM to model them (Lakshmanan et al, 2015). In contrast, our work models trial-to-trial variability as shared, rather than unique, to each neuron.

**Table 2:**
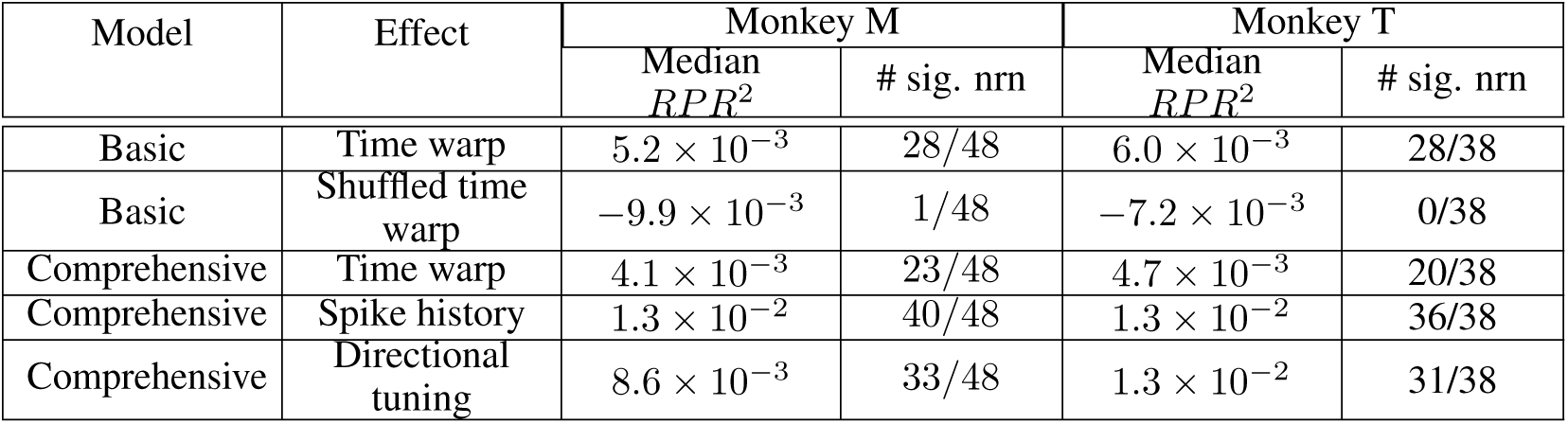
Comparison of effect sizes. For each neuron, the effect size of a given model component (e.g., time-warp variability) was calculated as the Relative Pseudo-*R*^2^ (*RPR*^2^) between the model with all components and the model lacking the component in question. For each effect, this table presents the median effect size (in terms of *RPR*^2^) across all neurons, as well as the number of individual neurons for which that effect was statistically significant (*p* < .05, bootstrap test). ‘Basic model’ refers to the simpler LNP+DTW model, which includes spatial tuning and time-warp variability components. ‘Comprehensive model’ refers to the model which additionally includes reach-specific kinematic (x-velocity, y-velocity, x-acceleration, y-acceleration, speed) and spike history components. The shuffled time warp analysis examined whether the time warp parameters were similar across trials (see Results and Methods for full description).

Our study builds on this previous work to formalize an alternative model of trial-to-trial temporal variability, and to quantify its magnitude. Importantly, it combines the GLM with DTW into a joint model that can be efficiently fit to data and benefits from both a model of spiking variability and a model of shared temporal variability.

Another recent study used a similar alignment-matrix approach based upon Mixed Pair Hidden Markov Models to study spike jitter (Kollmorgen and Hahnloser, 2014). Using simulations, Kollmorgen et al found that a model incorporating jitter allowed better receptive field estimation. They also applied their model to single-neuron data and found modest improvements in fit. Our approach based upon Dynamic Time Warping is theoretically similar, but our model applies to a *population* of simultaneously recorded neurons rather than a single neuron. This is important for a number of reasons. First, our approach forms a bridge between models of spike jitter (Kollmorgen and Hahnloser, 2014; Aldworth et al, 2011, 2005; Gollisch, 2006) and latent variable models of shared temporal variability (Vidne et al, 2012; Goris et al, 2014; Chase et al, 2010; Lawhern et al, 2010; Pfau et al, 2013; Lawhern et al, 2010). Second, modeling a population of neurons makes it possible to perform a rigorous cross-validation procedure not possible in their work: we are able to fit the trial-specific time-warp path on one subset of neurons, and test it on a separate, held-out subset; this allows for more accurate estimation of effect size. Third, using simulations, we found that time-warp path estimates are quite poor using small numbers of neurons, and by extension, using a single neuron (see Supplemental Results, Fig. S1). We also compared the effect size of time-warp variability to those of spike history and directional tuning, key effects in real data. This is important for contextualizing the degree of LNP+DTW model improvement. Their study has the benefit, however, of *fitting* the DTW-like parameters by integrating over all possible alignment paths; we specify the DTW parameters as priors. This may mean their model provides somewhat better fits, but we find that the values of these DTW parameters become less important with increasing numbers of neurons (see Supplemental Results, Fig. S1). While the two studies use similar alignment-matrix approaches, we believe that our study builds upon their excellent work in a number of important ways.

There are always multiple sources of variability present in spike trains. Other models of non-Poisson variability, such as latent variable models (Vidne et al, 2012; Yu et al, 2009; Goris et al, 2014; Pfau et al, 2013; Buesing et al, 2012; Shenoy et al, 2013), are likely to offer an alternative description of some of the variability we describe here. But the same is likely true for any study of neural variability (Gold and Shadlen, 2007; Latimer et al, 2015); models and interpretations in neuroscience are often formulated at different levels of mechanistic abstraction. In any case, by preserving easily interpretable models of single neurons, we believe that our model of time-warp variability is conceptually different from LVMs. These sources of variability could, in the future, be combined in a single model. For example, a gain parameter (as in Goris et al, 2014) may explain some of the same variability as does our model (see Supplemental Results for a similar analysis, Fig. S3). Lastly, some of the effect of time warp could be attributed to not knowing when each movement “starts”. Here, we use a speed threshold to label movement onset, but this is almost certainly an imperfect model. To explore this issue, we examined a variety of different threshold speeds and found the effect of time warp in all cases (see Supplemental Results, Fig. S4). Modeling time-warp variability is one way to cope with and quantify this type of ignorance. Nevertheless, future work should combine the full breadth of known sources of variability into a single model.

A limitation of this study is that we were not able to model the sensory component of our task, as target presentation times were not recorded precisely enough. PMd is known to encode both sensory and motor variables (Cisek and Kalaska, 2005), meaning that some time-warp variability may instead be unmodeled sensory variability (e.g., variable reaction times). However, we believe it is unlikely that this property would explain away *all* time-warp variability. PMd includes cells that span a variety of response types, not just those that respond to target presentation (Weinrich et al, 1984; Crammond and Kalaska, 2000; Cisek and Kalaska, 2004; Churchland and Shenoy, 2007). We believe that internal factors such as attention and arousal, in addition to reaction time, are likely to increase time-warp variability. Future work should, nonetheless, strive to build more complete models than we present here.

Our study emphasizes that the temporal dynamics of higher-order brain activity are quite complex. Realistically, no two movements in the natural world are exactly the same, in terms of planning or execution. But whereas movement execution is measurable, movement planning represents an internal state that has to be inferred. Furthermore, higher-order brain areas involved in planning and deliberation are likely to be both multi-modal and highly processed compared to lower-order brain areas. It should not be surprising, therefore, that we find effects like time warping, especially when considering less-constrained tasks.

Future work could extend our approach in multiple ways. Here we do not seek an explanation for the cause of the time warps. It may be possible to identify the factors that predict time warp on a trial-by-trial basis, whether they be related to behavior, or to circuit mechanisms (Guo et al, 2017). It may also be possible to fit more complicated time-warp models, e.g., with different time-warp paths for different types of covariates, or with different time-warp paths for different subpopulations of neurons. Lastly, time-warp models could be combined with other types of models, e.g., latent variable models, to delineate the sources of variability in spike trains.

We believe that time warping is a general phenomenon in the brain. Temporal jitter has been reported throughout the nervous system (Siegel et al, 2015). We expect that every step away from the periphery is likely to add both *fixed* time delays, as well as *variable* time delays arising from a number of sources: recurrent connections, attractor dynamics, or variability in planning among others. This may mean that high-level brain areas are effectively decoupled from the periphery. As such, we believe that the need to model time warping may be universal in neuroscience.

## Methods

### Behavioral tasks

Two monkeys (Monkey M, MM; Monkey T, MT) performed a reaching task in which they controlled a computer cursor using arm movements (Fig. 6). The monkey was seated in a primate chair while operating a two-link planar manipulandum. Hand movements were constrained to a horizontal plane within a workspace of 20 cm x 20 cm. In this task, an on-screen visual cue (2 cm square) specified the target location for each reach, and after making a series of four correct reaches to the targets, the monkey was given a liquid reward. The location of the target was pseudo-randomly chosen to be located within an annulus (radius = 5-15 cm, angle = 360 degrees) centered on the current target.

The timing of target presentation was different for the first reach of each trial (beginning of series; made from rest) than for reaches two through four (mid-series). For the first reach, the target was presented and the monkey was allowed to move without an instructed delay period. For reaches two through four, the monkey initiated the next target by holding the cursor briefly within a 2 cm x 2 cm box centered on the current target. Upon reaching the current target, the next target was triggered (NB: not displayed) 100 ms later. The time of next target appearance was not precisely recorded, but was determined with later experiments to appear on average 96 ms after being triggered. Thus, on average, the next target appeared approximately 200 ms after the monkey reached the current target. Additionally, there was an imposed hold period of 100 ms that began at the same time as the triggering of the next target. Thus, there was a 200 ms time interval between when the monkey reached the current target and when it was allowed to initiate the next movement. This brief hold period had the effect of forcing the monkeys to decelerate as they approached the target. In practice, the task led to a series of relatively smooth arm movements of variable distance.

All surgical and experimental procedures were consistent with the guide for the care and use of laboratory animals and approved by the institutional animal care and use committee of Northwestern University.

### Data acquisition and processing

MM and MT were both implanted with 100-electrode arrays (Blackrock Microsystems, Salt Lake City, UT) in dorsal premotor cortex (PMd). We excluded units with low firing rates (< 2 spikes/sec), as well as units having high correlations with other units (to guard against duplicate units resulting from crosstalk between electrodes and imperfect spike-sorting; excluded units with *r* > 0.4). MM yielded 95 well-isolated units, of which 47 were excluded, leaving 48 for analysis. MT yielded 49 well-isolated units, of which 11 were excluded, leaving 38 for analysis. These processing decisions were made independent of the analysis results.

Only successful reaches were included in the analysis (those that were held at the target for at least 100 ms). In this study, we built models of movement encoding, and to do this we defined movement initiation using a speed threshold. Here we used 8 cm/s, but in order to test for sensitivity to this parameter, we fit the model using other values for the threshold (see Supplementary Results). To distinguish drift from discrete arm movements, we excluded arm movements not meeting a minimum peak speed of 12 cm/s (chosen by eye, but independent of neural recordings and analysis results). We also excluded reaches with multiple speed peaks to in order to focus on simple reaches.

### Neural data analysis and modeling

#### General

In this study, we modeled temporal variability in spike trains that differs from trial to trial but that is shared across neurons (time-warp variability). To do this, we combined the Linear-Nonlinear-Poisson (LNP) model of neural activity, and Dynamic Time Warping (DTW). We proposed a generative model (outlined in the Results section), tested the model with simulations, and then applied the model to PMd data. Model code is available on Github (https://github.com/pnlawlor/LNTWP), and data will be available at CRCNS. Here we describe the model in further detail.

#### Linear-Nonlinear-Poisson (LNP) Model: General

We modeled extracellularly recorded spiking activity using the Linear-Nonlinear-Poisson (LNP) model (also known as the Poisson Generalized Linear Model). The LNP treats the spike train as a binned inhomogeneous Poisson process (i.e., with a time-varying firing rate). To model this non-negative firing rate, explanatory features are linearly combined and then passed through an exponential nonlinearity (the Poisson inverse link function) (Fig. 2; described in Results). The number of spikes in each 10 ms time bin is then drawn randomly from a Poisson distribution with the mean given by the estimated firing rate in that bin.

To apply our model to real data, we analyzed PMd activity recorded during a reaching task. We therefore designed LNP covariates so as to accurately model its activity. Generally speaking, PMd neurons are thought to be involved in motor planning (Cisek and Kalaska, 2005). An individual neuron’s activity is often related to reaches towards a particular direction, and occurs with a roughly stereotyped temporal response. We therefore modeled both the spatial and temporal aspects of neuronal activity. Details of the parameterization and fitting procedure are provided below.

#### LNP: Spatial tuning parameterization

To model the spatial component of PMd activity, we parameterized space using polar coordinates (angle and eccentricity). We used cosine tuning for the angular coordinate, and flat tuning for the eccentric coordinate (no dependence on reach length). We chose flat eccentric tuning in order to simplify the model, although visual examination of PSTHs suggested that this was a reasonable simplification (data not shown).

#### LNP: Temporal response parameterization

Rather than model all data recorded during these experiments, we selected a fixed temporal window surrounding each reaching movement. (400 ms before reach onset, 200 ms after reach onset; 10 ms bins). This interval is large enough to capture the movement-related activity associated with each reach.

The temporal responses of PMd neurons are heterogeneous, so we allowed for substantial variability in their shapes. To do this, we used a variety of temporal features that spanned the window of interest. More specifically, we convolved the spatial tuning features of each reach with a set of temporal basis functions (raised cosines) of multiple widths and temporal offsets. Because we sought to build a model of movement, these temporal basis functions were aligned to movement onset. This broad set of temporal basis functions allowed us to model a variety of temporal responses.

#### LNP: Basic and Comprehensive models

The basic LNP model of movement activity in PMd is as follows (Fig. 11):

1. Baseline firing rate: We used a scalar term, *θ*_0_, to model a constant baseline firing rate.
2. Untuned temporal responses: Neurons may have reach-related responses that do not depend on the direction of the reach (Fernandes et al, 2013). We accounted for this possibility by using separate untuned temporal responses aligned to reach onset. Intuitively, these temporal basis functions make it possible to explain spiking of variable duration and temporal offset with respect to reach (Fig. 11). The untuned reach-related response is given by

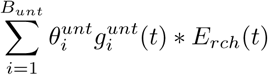 The 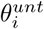 are the free parameters, 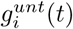 are the temporal basis functions (raised cosine functions, 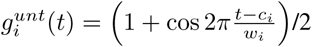; widths = {75,100,150,200} ms; offsets were chosen separately for each width and were designed to span the temporal window), and *E*_*rch*_(*t*) is an “event function” that specifies the time of reach onset (*E*_*rch*_ = 1 at reach onset, *E*_*rch*_ = 0 otherwise). We used 20 temporal basis functions in total (*B*_*unt*_ = 20), and thus the untuned temporal response was specified by 20 total parameters.
3. Spatiotemporal reach tuning: We modeled the neural activity around a reach event as a function of the upcoming reach direction. To do this, we constructed the movement covariates as the sine and cosine projections of the upcoming reach direction. We did not incorporate previous knowledge of the neuron’s preferred direction (PD), Ψ (reach angles were defined as the angle formed by the vector connecting the starting position and ending position of the reach; Fig. 11). Rather, the neuron’s PD, Ψ*, can be inferred from the data by using the fitted parameters and trigonometric identities:

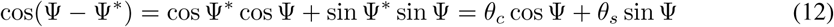 This yields 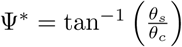. The spatiotemporal reach tuning portion of the model is given by

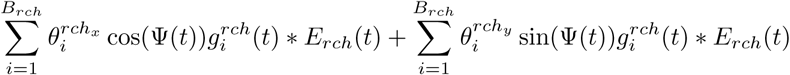 The functions 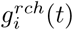 and *E*_*rch*_(*t*) are identical to those of the untuned model. The spatiotemporal reach response was therefore specified by 40 total parameters (*B*_*rch*_ = 20; 2 reach axes). Note that our model is not specified as a bilinear model as in Fernandes et al (2013); Ramkumar et al (2016), i.e., whose spatiotemporal response is multiplicatively separable. Although a bilinear model is more easily interpreted (one preferred direction for all temporal basis functions), it is more complicated to fit because it is not linear in the model parameters. The model used here is more flexible and easier to fit (linear in parameters), but the parameters are more difficult to interpret (potentially a different preferred direction associated with each temporal basis function). We have chosen this model because of its computational simplicity and because we are not interested in making inferences about PDs. We believe that this choice is unlikely to affect our results; if anything, it allows for more flexibility in the LNP model than the bilinear model, decreasing the potential explanatory power of the DTW model component. The following model components were included in the comprehensive LNP model in addition to components 1-3 in the basic model:
4. Kinematic tuning: We modeled the effects of kinematic variables on neural activity, as this is one possible source of trial-to-trial variability. In particular, we used the sine and cosine components of instantaneous velocity and acceleration, as well as instantaneous speed. This portion of the model is given by:

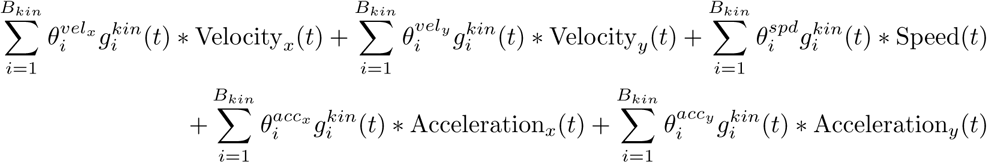 Here, e.g., Velocity_*x*_(*t*) refers to the instantaneous reach velocity in the *x* direction. The 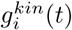 were again temporal basis functions, and we used two for each kinematic variable (raised cosine functions; width = 75 ms, offset = {–150,0} ms with respect to instantaneous kinematics). These convolution operations had the effects of a) smoothing the kinematic variables and b) allowing them to be offset with respect to neural activity. Modeling the offset is important because neural activity related to a reach likely precedes the reach. Thus, if instantaneous velocity is intended to explain the neural activity related to that reach, it must be shifted backwards in time. The kinematic tuning portion of the model was specified by 10 parameters (*B*_*kin*_ = 2; 5 kinematic variables considered).
5. To explain additional variability, we included spike history terms. We also asked whether spike history terms explained away the effects of time-warp variability. To model the effect of spike history, we convolved the spike train, *y*(*t*), with 5 raised-cosine temporal basis functions, 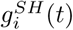 (raised cosine functions; widths = {30,40, 80,140,270} ms, and corresponding offsets = {10, 30,40, 70,120} ms from spike time) (Pillow et al, 2008). This portion of the model is given by

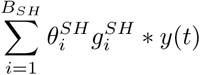 The spike history portion of the model was specified by 5 parameters (*B*_*SH*_ = 5).

#### DTW: General and model fitting

To correct the LNP model predictions for time-warp variability, we used the Dynamic Time Warping (DTW) algorithm. Given two signals, DTW finds the optimal transformation (consisting of multiple, piecewise offset and scaling operations) that aligns them (Fig. 3). The computational complexity of finding the globally optimal solution is *O*(*T*^2^) per trial of duration *T* time bins using dynamic programming (see Supplemental Methods for intuition) (Berndt and Clifford, 1994; Sakoe and Chiba, 1978).

#### DTW: Priors

The DTW algorithm finds the optimal alignment transformation between two signals by calculating a path through signal_1_ × signal_2_ space (Fig. 3). Because the signals are discrete, this path consists of discrete steps. The nature of these steps can be tweaked using a small number of free parameters in the DTW algorithm (Eqn. 6, second term). These parameters can be chosen to prohibit certain types of steps as well as to adjust the relative prior probabilities of different types of steps.

In the case of basic DTW, the following types of steps are allowed and equally likely: up 1, over 1, and up 1 and over 1 (diagonal). Here, however, we use a slightly different set of parameters in order to accommodate our generative model for time-warp variability in spike trains. We frame the estimation problem as follows: for every time *t* in spike train time, we would like to estimate the time *τ* in LNP model-prediction time that is most consistent with the observed spikes at (subject to continuity and forward-progression in time; Eqn. 9). In other words, we view the spike trains as observed (fixed) data, and would like to warp the LNP model predictions to best fit the spike trains. Thus, we chose step parameters to avoid warping the spike trains themselves (i.e., no spikes are dropped or added; the model’s predictions are warped to be most compatible with the spikes). This amounted to requiring that spike-train time, *t*, increment by exactly one bin with each step. We allowed the model time, *τ*, to increment by 0 (stretch), 1 (no time warp), or 2 (skip/compression) with each step (Fig. 4). These parameters made it possible to warp the LNP model predictions to the spike trains as well as to accommodate our generative model.

#### LNP + DTW: General

To model time-warp variability, we combined the LNP model with DTW into a statistical model. The goal of this model was to use the LNP model to capture the average responses of neurons, and then to use DTW to correct the LNP model on a trial-by-trial basis (Fig. 4). We are ultimately interested in modeling the number of spikes 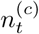 for each cell *c* that occur in a given time bin *t* of duration *dt*. We assume that 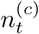 is Poisson-distributed with a mean given by 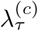, which reflects an underlying firing rate that varies in time. The mean, in turn, depends on the parameters of the standard LNP process, Θ^(*c*)^, as well as *X*, and the time-warp path, *τ*. The LNP parameters, Θ^(*c*)^, describe a neuron’s dependence on external variables (e.g., reach direction), *X* (dependence on internal variables such as spike history can also be included in the model; see Control Analyses subsection below). Each neuron *c* is described by its own set of parameters Θ^(*c*)^ which are assumed to be fixed across trials and independent of other neurons. The time-warp parameter, *τ* = {*τ*_1_, *τ*_2_,…,*τ*_T_}, is modeled as a nonlinear mapping between the LNP model time, *τ*, and spike time, *t*. We assume that a single *τ*^(*j*)^ is shared across neurons, and varies from trial to trial (with trial parameterized by *j*). We model the covariates *X* as parameterized in time by *τ*, and the spikes as parameterized in time by *t*. To summarize, we model spiking as dependent upon external covariates that are time warped from trial to trial and shared across neurons.

#### LNP + DTW: Model fitting

To fit the combined LNP+DTW model, we used an EM-like approach in which we alternated between fitting the LNP and fitting the DTW portions of the model. I.e., we first fit the LNP; we then fit DTW while holding the LNP parameters constant; we then re-fit the LNP parameters while holding the DTW parameters constant, and so on. We chose this approach because it is difficult to optimize for both the Θ^(*c*)^ and *τ*^(*j*)^ directly. This would amount to solving

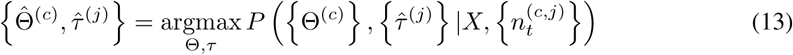

See Supplementary Methods for the full derivation of the likelihood equations and inference algorithm.

### Control analyses and other model components

To rule out the possibility that the improvement in model fit of the LNP+DTW algorithm was spurious, we performed a number of control analyses.

First, we asked if the addition of DTW simply improved the estimates of neurons’ average temporal responses. Although the temporal basis function set we used allowed for considerable variability, it might still not describe all possible temporal responses adequately. It is possible, then, that DTW could be performing some kind of generic operation that better shapes the temporal aspects of the LNP model predictions (e.g., by smoothing). If this were true, the calculated time warp should be similar across trials, and thus interchangeable across trials. To test this, we performed a shuffle analysis in which we calculated time warps on the correct trials, but then applied them to other trials at random (i.e., corrected the LNP models of other trials at random). To ensure that this randomization procedure was applied fairly, we only shuffled between trials within the same 45 degree range of reach directions.

Next, we asked if other factors could explain away the effect of time warp. The LNP+DTW model seeks to explain “bouts” of spiking activity that are shared across neurons but differ from trial to trial. One possibility is that these bouts of activity are better explained by spike history terms than time warping. For example, a spike history kernel that favored bursting could possibly explain a bout of spiking activity. Another possibility is that trial-to-trial differences in kinematics could explain trial-to-trial differences in spike trains.

To test these possibilities, we fit a more comprehensive model designed to allow these factors to explain away the effect of time warp. In addition to the standard reach covariates (see basic LNP model and Fig. 11), we included spike history covariates and kinematic covariates (instantaneous velocity, speed, and acceleration) in the LNP portion of the model. Time warping was therefore allowed to explain only additional variability above and beyond these factors. See comprehensive LNP model for the parameterization of these model components.

### Model testing and comparison

To avoid overfitting, we used two types of cross validation to test our model (Fig. 12). To avoid overfitting the LNP model, we used 2-fold, trial-wise cross validation for each neuron (train on half of the trials, test on the held-out half; alternating trials). To avoid overfitting the DTW part of the model, we used 10-fold “leave-neurons-out” cross validation. To do this, we trained the DTW model on one subset of neurons, and tested it on the held-out subset. These cross-validation procedures helped to ensure that no part of the model was overfit and would thus generalize better. Importantly, the fact that similar time-warp paths are found in distinct subsets of neurons supports the view that the temporal variability is indeed shared across neurons.

**Figure 12:**
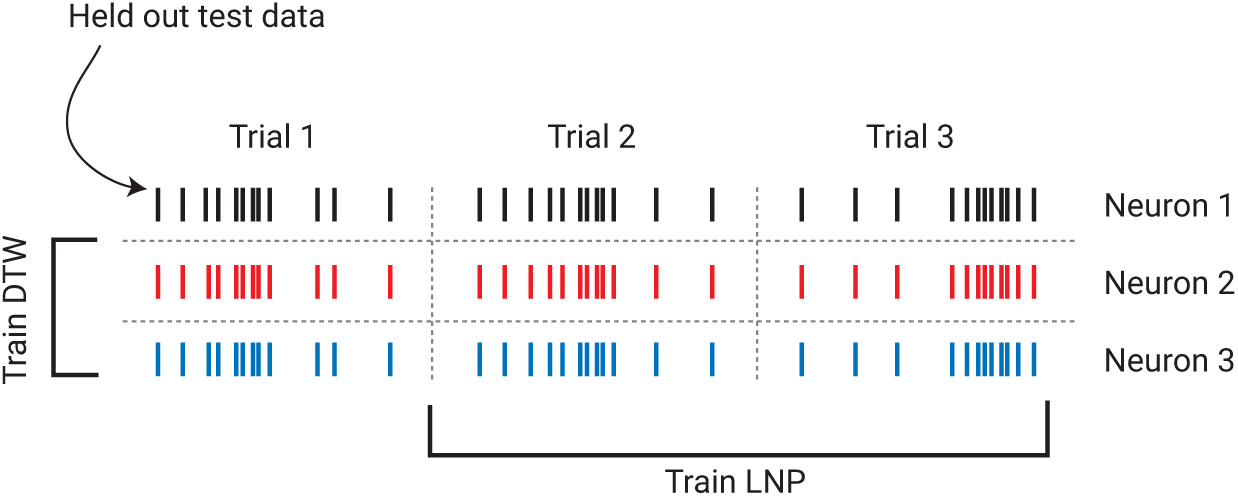
Cross validation procedures. We used two forms of cross validation to avoid over-fitting the model. To avoid overfitting the LNP model, we used trial-wise cross validation (train on one subset of trials, test on another). To avoid overfitting the DTW model, we used neuron-wise cross validation (train on one subset of neurons, test on another). This made it possible to assess model performance on truly naïve test data.

To assess model fit we used a metric called Pseudo-*R*^2^, a generalization of the standard *R*^2^ to target variables other than the Gaussian (here, to the Poisson distribution). It maps a model’s log likelihood to a value between 0 and 1 by scaling it to be between that of the simplest possible model (a constant/baseline model) and the most complex possible model (a “saturated” model consisting of perfect predictions). It is important to note that the absolute values of this metric are not comparable to (are much smaller in magnitude than) typical *R*^2^ for normally distributed variables. Where possible, we have provided reference values. Furthermore, all log likelihoods and accuracy metrics were computed on held-out test-set data.

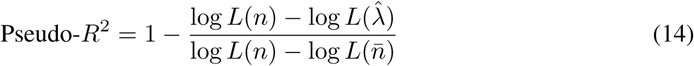

Here, log *L* refers to the log likelihood of the model. 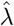 refers to the model in question; *n* refers to the saturated model (perfect predictions); 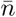 refers to the baseline model consisting only of the mean of the data. To compare two separate models, we used a related metric called Relative Pseudo-*R*^2^. Instead of comparing the model in question (e.g., model 2) with a baseline model, it compares it with another model (model 1). This metric, however, can take on negative values if model 1 is better than model 2. Relative Pseudo-*R*^2^ is often used to compare a more complex model with a simpler, nested model. In this case, Relative Pseudo-*R*^2^ would reflect the marginal improvement gained by model 2’s additional complexity. In this study, for example, we calculate the Relative Pseudo-*R*^2^ between the LNP model and the LNP+DTW model.

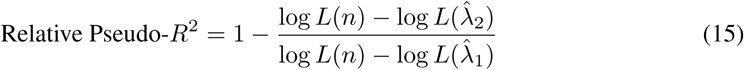

Here, 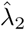 refers to the mean value predicted by model 2, the more complex model; 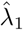 refers to the mean value predicted by model 1, the simpler or nested model. The other variables are the same as in Eqn. 14.

### Simulations

Because we proposed a new type of model, we sought to characterize its behavior using simulations before applying it to real data. To do this, we simulated spike trains using the generative model, and then attempted to recover the model parameters without knowledge of the simulations. See Supplementary Methods for the full methodology of the simulation design.

